# Inhibitory synaptic transmission is impaired at higher extracellular Ca^2+^ concentrations in *Scn1a*^+/−^ mouse model of Dravet syndrome

**DOI:** 10.1101/2021.02.04.429687

**Authors:** Kouya Uchino, Hiroyuki Kawano, Yasuyoshi Tanaka, Yuna Adaniya, Ai Asahara, Masanobu Deshimaru, Kaori Kubota, Takuya Watanabe, Shutaro Katsurabayashi, Katsunori Iwasaki, Shinichi Hirose

## Abstract

Dravet syndrome (DS) is an intractable form of childhood epilepsy that occurs in infancy. More than 80% of all patients have a heterozygous abnormality in the *SCN1A* gene, which encodes a subunit of Na^+^ channels in the brain. However, the detailed pathogenesis of DS remains unclear. This study investigated the synaptic pathogenesis of this disease in terms of excitatory/inhibitory balance using a mouse model of DS. We show that excitatory postsynaptic currents were similar between *Scn1a* knock-in neurons (*Scn1a*^+/–^ neurons) and wild-type neurons, but inhibitory postsynaptic currents were significantly lower in *Scn1a*^+/–^ neurons. Moreover, both the vesicular release probability and the number of inhibitory synapses were significantly lower in *Scn1a*^+/–^ neurons compared with wild-type neurons. There was no proportional increase in inhibitory postsynaptic current amplitude in response to increased extracellular Ca^2+^ concentrations. Our study revealed that the number of inhibitory synapses is significantly reduced in *Scn1a*^+/–^ neurons, while the sensitivity of inhibitory synapses to extracellular Ca^2+^ concentrations is markedly increased. These data suggest that Ca^2+^ tethering in inhibitory nerve terminals may be disturbed following the synaptic burst, likely leading to epileptic symptoms.

## INTRODUCTION

Epilepsy is a pervasive and severe neurological disorder that affects 0.5%–1% of the population, predominantly in childhood^1^. Epilepsy patients are generally treated with antiepileptic drugs, but about 30% of patients have refractory epilepsy that does not respond to such treatment^2^. The molecular mechanisms underlying the onset of refractory epilepsy remain mostly unknown. Therefore, epilepsy is one of the most challenging diseases for developing fundamental therapies and drugs. In addition, pediatric patients with intractable epilepsy may develop complications, such as developmental disorders and mental retardation, leading to anxiety about their social lives even in adulthood^3,4^.

We have previously identified more than 100 genetic abnormalities in Na^+^ channels via the genetic analysis of patients with intractable epilepsy resulting in mental retardation^5,6^. Dravet syndrome (DS) is a severe form of epilepsy that develops in infancy and causes frequent seizures, leading to severe encephalopathy and developmental disorders. Heterozygous mutations in *SCN1A*, the gene encoding Na_v1.1_ (the α subunit of voltage-gated Na^+^ channels), have been identified in more than 80% of patients with DS. However, it remains unclear how this gene mutation causes intractable epilepsy. Recently, knock-in, knock-out, and conditional knock-out mice with heterozygous mutations in the *Scn1a* gene have been reported as mouse models of DS^7,8,9,10^. In parallel with the development of mouse models, we established induced pluripotent stem cells from a DS patient with a heterozygous nonsense mutation in the *SCN1A* gene^11,12^. We reported an apparent fragile mutation in the maintenance of action potentials during sustained depolarizing stimulation of γ-aminobutyric acid (GABA)ergic neurons. The common mechanism of these aforementioned studies was that *SCN1A* mutations result in a loss of inhibitory neural function. This underlying mechanism indicates that excitatory neural function is increased in the brain, making it highly susceptible to epilepsy because of the suppression of inhibitory neural function in the central nervous network.

In the brain, neurons connect via synapses and transmit information in a fine-tuned manner to maintain central homeostasis. During epileptogenesis, the excitatory/inhibitory (E/I) balance of the multicellular neural network is transiently or persistently disrupted, resulting in overexcited synaptic activity^13,14^. Therefore, understanding epilepsy as a “synaptic pathology” and elucidating synaptic recovery mechanisms may help to develop fundamental epilepsy therapies and aid in drug discovery.

## RESULTS

### Establishment of Scn1a-targeted mice

The targeting vector consisted of the neomycin-resistance cassette, of which the 5’ and 3’ ends were connected to recombination arms derived from intron 4 to intron 7 (left arm) and intron 12 to intron 16 (right arm) of the mouse *Scn1a* gene, respectively (Figure 1a). After this was introduced into mouse ES cells, 156 neomycin-resistant clones were isolated and screened for targeted homologous recombination by PCR. Using primers corresponding to the neo gene and intron 16, a 5.5-kb fragment that would emerge only from recombinant alleles was observed in three clones. These candidates were further analyzed by Southern hybridization. The full length of the gel blot is shown in the Supplementary Information (Figure S1), and a cropped image from the full gel blot is shown in Figure 1c. Two of them showed positive signals at 7.8 and 6.5 kb, which corresponded to the wild-type and targeted alleles, respectively (Figure 1b, c). These two clones were regarded as having desirable recombination and were used to establish the mutant mouse, in which exon 8 to exon 12 of the *Scn1a* gene was heterogeneously replaced by the neomycin resistance cassette (*Scn1a*^+/–^ mouse). On establishment and after every mating, mouse specimens with recombinant alleles were identified by PCR amplification of a 634-bp DNA fragment spanning from intron 7 to the 5’ FRT element.

**Figure 1.**
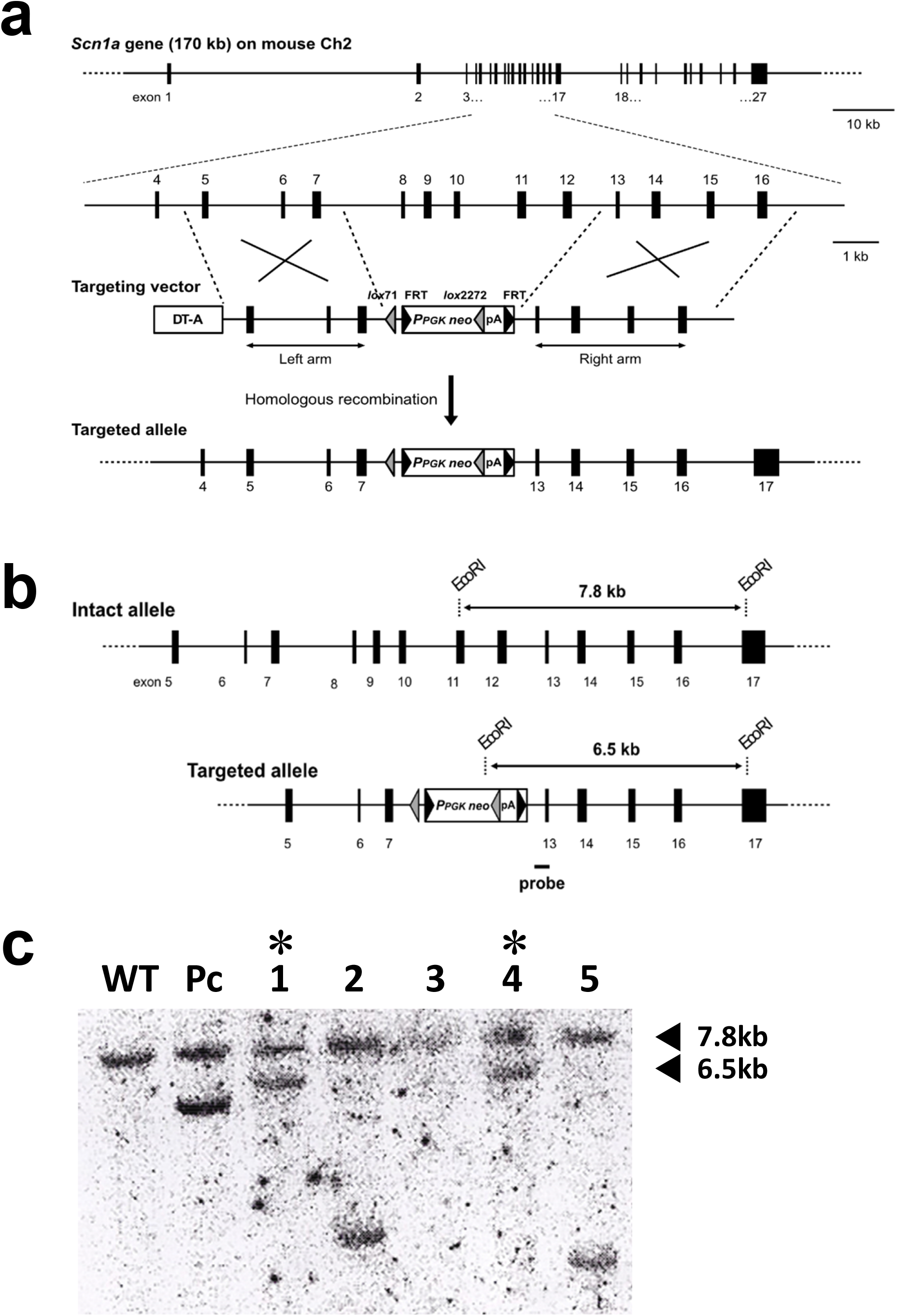
Targeting of the *Scn1a* gene in mouse embryonic stem cells. (**a**) Strategy for *Scn1a* targeting. Upper, overall schematic view of the mouse *Scn1a* gene, and a magnified view containing the targeted region from exons 8 to 12. Exons are shown as numbered solid boxes, and introns are shown as horizontal lines. Middle, the organization of the targeting vector. More detail is indicated in the Materials and Methods. Shaded and solid triangles indicate the lox and flippase recognition target sequence (FRT) elements, respectively. Regions to be recombined between genomic *Scn1a* and the targeting vector are indicated by large Xs. Bottom, the targeted *Scn1a* allele that resulted from desirable homologous recombination. (**b**) Comparison of the restriction patterns of wild-type and targeted *Scn1a* alleles in Southern analysis. The position of the hybridization probe is indicated by a solid bar. (**c**) A result of Southern hybridization for seven candidate clones. For samples #1 and #4, both the 7.8- and 6.5-kb fragments were detected, which were derived from the wild-type and homologously recombined *Scn1a* alleles.

### Excitatory synaptic transmission of *Scn1a*^+/–^ neurons

In the epileptic brain, changes in the E/I balance are exhibited as enhanced excitatory synaptic transmission and/or reduced inhibitory synaptic transmission, and vice versa. We first measured excitatory synaptic transmission. The EPSC amplitudes were not significantly different between WT and *Scn1a*^+/–^ neurons (Figure 2a, b). The mEPSC amplitudes were also not significantly different between WT and *Scn1a*^+/–^ neurons (Figure 2c, d). We then analyzed the frequency of mEPSCs to assess the effects on presynaptic machinery, and found no significant differences between WT and *Scn1a*^+/–^ neurons (Figure 2e). These results indicate that there are no changes in the excitatory synaptic transmission of *Scn1a*^+/–^ neurons.

**Figure 2.**
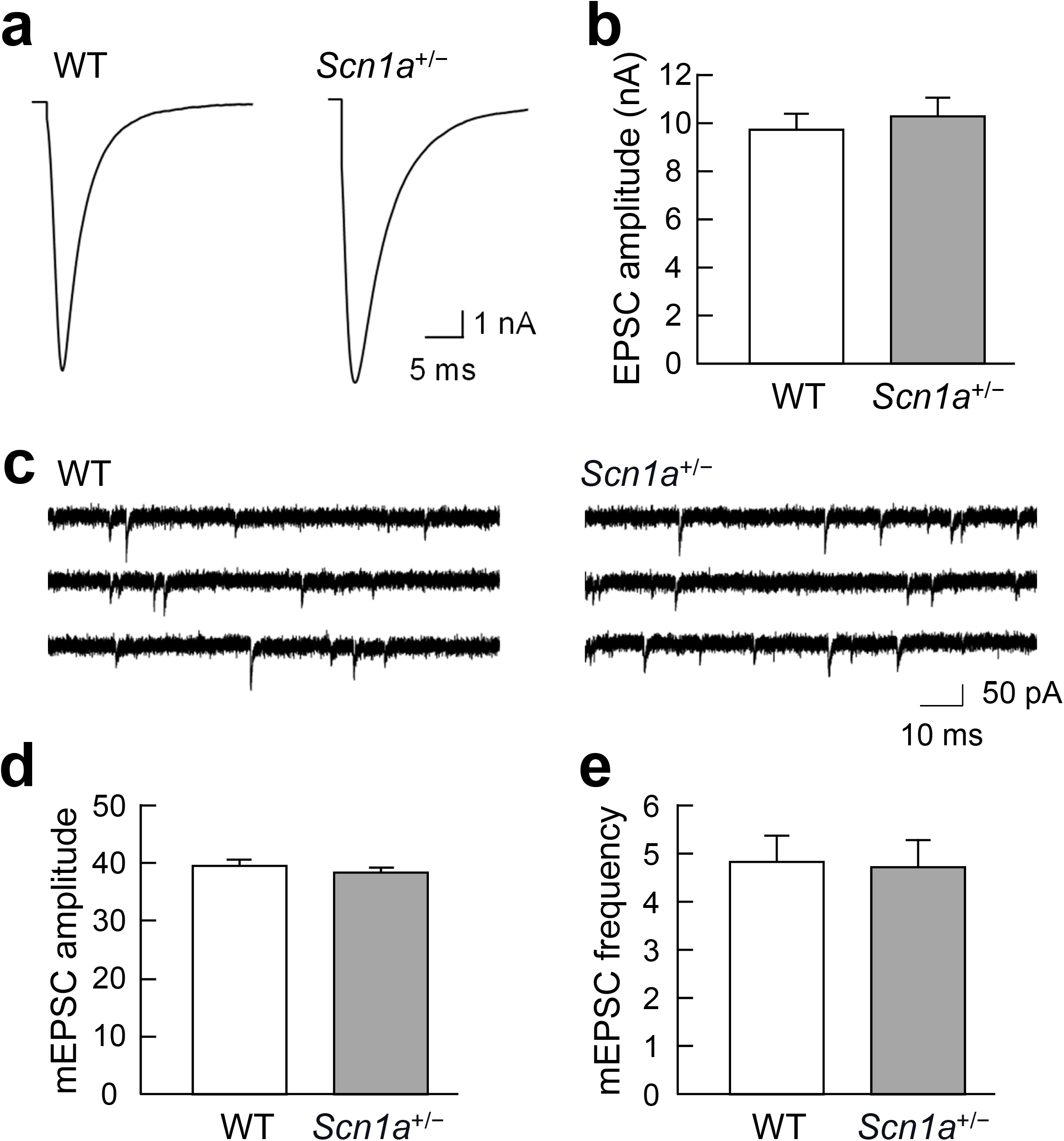
No changes in the excitatory synaptic transmission of *Scn1a*^+/–^ neurons. (**a**) Representative traces of evoked excitatory postsynaptic currents (EPSCs) recorded from a single autaptic hippocampal neuron of either wild-type (WT) or *Scn1a*^+/–^ mice. Depolarization artifacts caused by the generated action currents have been removed for clarity of presentation. (**b**) Average amplitudes of the evoked EPSCs in neurons of WT or *Scn1a*^+/–^ mice (WT: 9.65 ± 0.62 nA, *n* = 104; *Scn1a*^+/–^: 10.25 ± 0.69 nA, *n* = 102/*N* = 5 cultures). (**c**) Representative miniature EPSC (mEPSC) traces in neurons of either WT or *Scn1a*^+/–^ mice. (**d**) Average mEPSC amplitudes in neurons of either WT or *Scn1a*^+/–^ mice (WT: 39.47 ± 0.87 pA, *n* = 104; *Scn1a*^+/–^: 38.15 ± 0.71 pA, *n* = 102/*N* = 5 cultures). (**e**) Average mEPSC frequencies in neurons of either WT or *Scn1a*^+/–^ mice (WT: 4.82 ± 0.52 Hz, *n* = 104; *Scn1a*^+/–^: 4.70 ± 0.56 Hz, *n* = 102/*N* = 5 cultures). Data were obtained from the same autaptic neurons as in (**d**).

### Inhibitory synaptic transmission of *Scn1a*^+/–^ neurons

The absence of phenotypes in the excitatory synaptic transmission of *Scn1a*^+/–^ neurons (Figure 2) suggested that the inhibitory synaptic transmission of these neurons may be reduced, in terms of the E/I balance between excitatory and inhibitory synaptic transmission during epileptogenesis. Therefore, in the following experiment, we analyzed the inhibitory synaptic transmission of *Scn1a*^+/–^ neurons.

As predicted, IPSC amplitudes (i.e., inhibitory synaptic transmission elicited by action potentials) were significantly lower in *Scn1a*^+/–^ neurons compared with WT neurons (Figure 3a, b). In contrast, mIPSC amplitudes did not differ significantly between *Scn1a*^+/–^ and WT neurons (Figure 3c, d).

**Figure 3.**
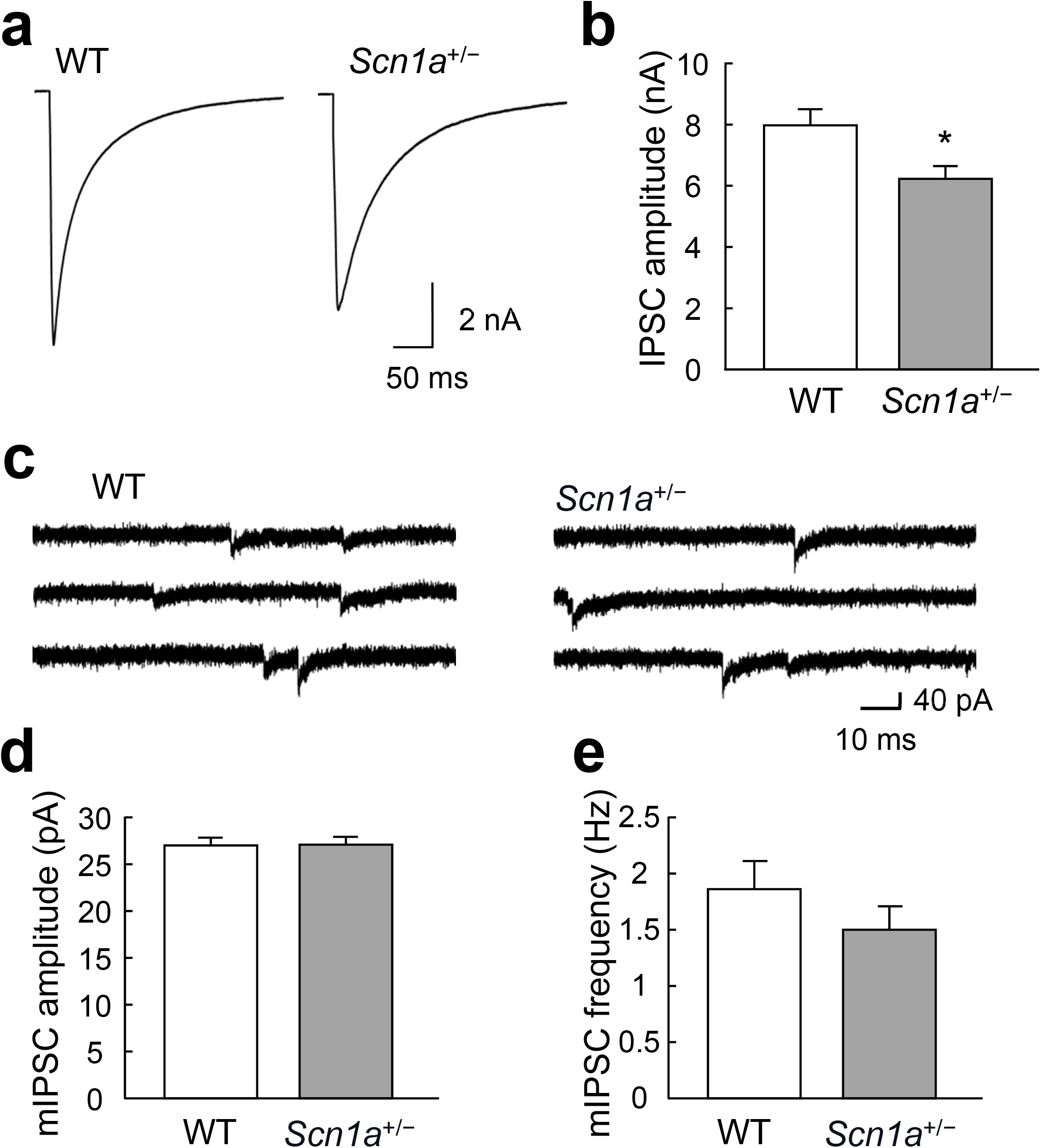
Changes in the inhibitory synaptic transmission of *Scn1a*^+/–^ neurons. (**a**) Representative traces of evoked inhibitory postsynaptic currents (IPSCs) recorded from a single autaptic hippocampal neuron of either wild-type (WT) or *Scn1a*^+/–^ mice. Depolarization artifacts caused by the generated action currents have been removed for clarity of presentation. (**b**) Average amplitudes of the evoked IPSCs in neurons of WT or *Scn1a*^+/–^ mice (WT: 7.94 ± 0.55 nA, *n* = 96; *Scn1a*^+/–^: 6.18 ± 0.42 nA, *n* = 98/*N* = 5 cultures, \**p* < 0.05). (**c**) Representative miniature IPSC (mIPSC) traces in neurons of either WT or *Scn1a*^+/–^ mice. (**d**) Average mIPSC amplitudes in neurons of either WT or *Scn1a*^+/–^ mice (WT: 26.94 ± 0.80 pA, *n* = 95; *Scn1a*^+/–^: 26.70 ± 0.86 pA, *n* = 96/*N* = 5 cultures). (**e**) Average mIPSC frequencies in neurons of either WT or *Scn1a*^+/–^ mice (WT: 1.85 ± 0.25 Hz, *n* = 95; *Scn1a*^+/–^: 1.50 ± 0.21 Hz, *n* = 96/*N* = 5 cultures). Data were obtained from the same autaptic neurons as in (**d**).

The finding that the amplitudes of IPSCs elicited by action potentials were reduced while the amplitudes of mIPSCs remained unchanged suggests that presynaptic function, rather than postsynaptic function, was affected in inhibitory synaptic transmission. To investigate presynaptic function, we analyzed the frequency of spontaneous mIPSCs. There were no significant differences in spontaneous mIPSC frequency between *Scn1a*^+/–^ and WT neurons (Figure 3e). These results suggest that inhibitory synaptic transmission in *Scn1a*^+/–^ neurons may be reduced by action potential-dependent mechanisms.

### Mechanisms of inhibitory synaptic transmission in *Scn1a*^+/–^ neurons

Because there was no significant decrease in the frequency of mIPSCs in *Scn1a*^+/–^ neurons, it remained unclear whether presynaptic function was impaired in these cells. We therefore measured the size of the RRP. There was no significant difference in RRP size between *Scn1a*^+/–^ and WT neurons (Figure 4a, b). However, the vesicular release probability (P_vr_) of synaptic vesicles was significantly lower in *Scn1a*^+/–^ neurons compared with WT neurons (Figure 4c).

**Figure 4.**
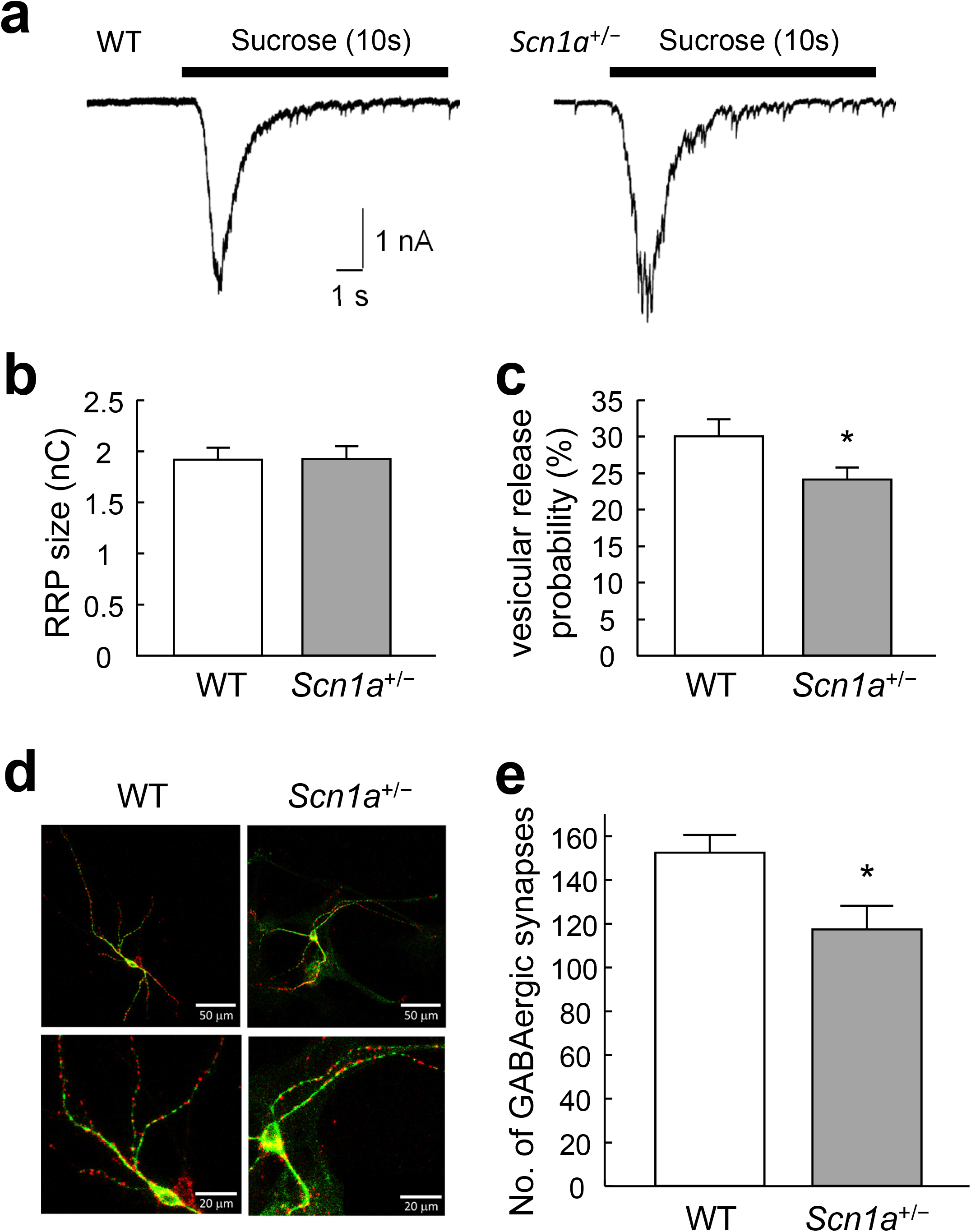
Changes in the number of inhibitory synapses in *Scn1a*^+/–^ neurons. (**a**) Representative traces of responses to 0.5 M hypertonic sucrose solution (10 s, black bar) in an autapse neuron of either wild-type (WT) or *Scn1a*^+/–^ mice. (**b**) The size of the readily releasable pool (RRP) of autaptic neurons of either WT or *Scn1a*^+/–^ mice (WT: 1.91 ± 0.12 nC, *n* = 91; *Scn1a*^+/–^: 1.92 ± 0.12 nC, *n* = 92/*N* = 5 cultures). (**c**) Vesicular release probability (P_vr_) in single autaptic neurons of either WT or *Scn1a*^+/–^ mice (WT: 29.99 ± 2.31%, *n* = 91; *Scn1a*^+/–^: 24.01 ± 1. 66%, *n* = 92/*N* = 5 cultures, \**p* < 0.05). (**d**) Representative images of autaptic neurons immunostained for the inhibitory nerve terminal marker vesicular GABA transporter (VGAT; red) and the dendritic marker microtubule-associated protein 2 (MAP2; green). Parts of the top row (scale bar 50 μm) are enlarged in the bottom row (scale bar 20 μm). (**e**) The number of VGAT–positive synaptic puncta in autaptic neurons of either WT or *Scn1a*^+/–^ mice (WT: 152.23 ± 7.91, *n* = 70, *Scn1a*^+/–^: 117 ± 10.61, *n* = 74/*N* = 8 cultures, \**p* < 0.05).

Reduced inhibitory synaptic transmission may also indicate that the number of inhibitory synapses is reduced. We therefore measured the numbers of inhibitory synapses in WT and *Scn1a*^+/–^ neurons using immunostaining. The number of GABAergic synapses was significantly lower in *Scn1a*^+/–^ neurons than in WT neurons (Figure 4d, e). These results indicate that the decreased inhibitory synaptic transmission in *Scn1a*^+/–^ neurons is likely caused by a combination of a lower P_vr_ and a lower number of inhibitory synapses.

### Mechanisms of depressed inhibitory synaptic vesicular release in *Scn1a*^+/–^ neurons

The generation of action potentials depolarizes nerve terminals, and the subsequent influx of extracellular Ca^2+^ into the nerve terminals causes activity-dependent neurotransmitter release. Thus, the lower P_vr_ in the inhibitory synaptic transmission of *Scn1a*^+/–^ neurons may be caused by a lower sensitivity to extracellular Ca^2+^ concentrations ([Ca^2+^]_o_). We therefore examined the changes in inhibitory synaptic transmission in response to higher or lower [Ca^2+^]_o_. There was no proportional increase in IPSCs at higher calcium concentrations (4 mM and 8 mM) in *Scn1a*^+/–^ neurons (Figure 5a). To further understand Ca^2+^ sensitivity, the relative amplitudes of evoked IPSCs were analyzed using Hill’s equation, with IPSC at 8 mM as a reference^15^. Compared with WT neurons, the graph of relative IPSC amplitude was shifted to the left in *Scn1a*^+/–^ neurons (Figure 5b), indicating that *Scn1a*^+/–^ neurons are more sensitive to [Ca^2+^]_o_. The K_d_ value calculated using Hill’s equation is a dissociation constant. Its reciprocal is a coupling constant, which can be approximated as the ion permeability in this case. In contrast, the Hill coefficient indicates cooperativity with ions. The K_d_ values were reduced to about one-third of WT values in *Scn1a*^+/–^ neurons (WT: 0.60 ± 0.05 mM, *Scn1a*^+/–^: 0.18 ± 0.10 mM), while Hill coefficients were higher compared with WT neurons (WT: 1.34 ± 0.15, *Scn1a*^+/–^: 2.03 ± 0.89). Thus, *Scn1a*^+/–^ neurons had higher affinity and cooperativity with [Ca^2+^]_o_ than WT neurons. These results indicate that the reduced inhibitory synaptic release in *Scn1a*^+/–^ neurons is not caused by low sensitivity to [Ca^2+^]_o_, but rather by high sensitivity to [Ca^2+^]_o_. In addition, the concentration of 2 mM [Ca^2+^]_o_ was hypothesized to be in the critical range for examining whether the vulnerability of the E/I balance was observed. When a larger number of data is recorded in the critical range, it is easier to detect a statistically significant difference. Indeed, Figure 3b shows the results from a relatively large number of recordings, which was large enough to detect a significant difference. In contrast, Figure 5a shows the results from a relatively small number of recordings, which did not show a significant difference between the two groups. Together, these findings indicate that specimen-to-specimen variation in the amplitudes of IPSCs may have masked changes in the release properties in the critical range (see the standard error bars in Figures 3b and 5a).

**Figure 5.**
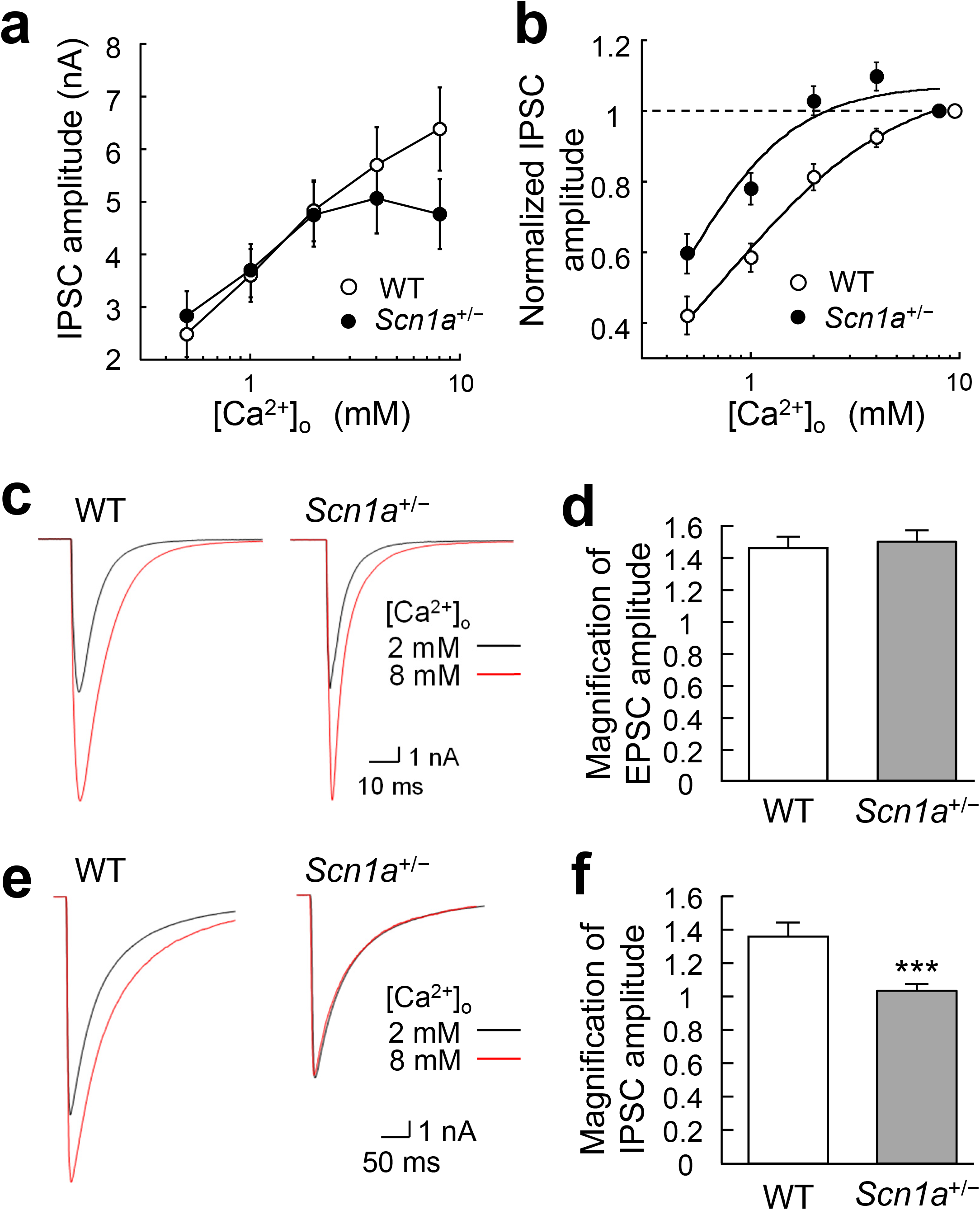
Changes in extracellular Ca^2+^ concentration sensitivity of inhibitory synapses in *Scn1a*^+/–^ neurons. (**a**) Inhibitory postsynaptic current (IPSC) amplitudes recorded in different extracellular [Ca^2+^]_o_ (0.5, 1, 2, 4, 8 mM) with 1 mM Mg^2+^ held constant in wild-type (WT) and *Scn1a*^+/–^ neurons (WT: *n* = 35, *Scn1a*^+/–^: *n* = 36/*N* = 9 cultures). (**b**) Dose-response curves and Hill fits for normalized IPSC amplitudes, recorded in different extracellular [Ca^2+^]_o_ (0.5, 1, 2, 4, 8 mM) with 1 mM Mg^2+^, held constant in WT and *Scn1a*^+/–^ neurons. IPSC amplitudes were normalized to the IPSC amplitude at 8 mM [Ca^2+^]_o_. Original data were obtained from the same cells as in (**a**). (**c**) Representative traces of evoked excitatory postsynaptic currents (EPSCs) recorded at 2 mM [Ca^2+^]_o_ (black line) and 8 mM [Ca^2+^]_o_ (red line) from neurons of either WT or *Scn1a*^+/–^ mice. Depolarization artifacts caused by the generated action currents have been removed for clarity of presentation. (**d**) Average magnification of evoked EPSCs at 8 mM [Ca^2+^]_o_ based on EPSC amplitudes at 2 mM [Ca^2+^]_o_ in neurons of WT or *Scn1a*^+/–^ mice (WT: 1.45 ± 0.07, *n* = 62, *Scn1a*^+/–^: 1.48 ± 0.07, *n* = 64/*N* = 11 cultures). (**e**) Representative traces of evoked IPSCs recorded at 2 mM [Ca^2+^]_o_ (black line) and 8 mM [Ca^2+^]_o_ (red line) from neurons of either WT or *Scn1a*^+/–^ mice. Depolarization artifacts caused by the generated action currents have been removed for clarity of presentation. (**f**) Average magnification of the evoked IPSCs at 8 mM [Ca^2+^]_o_ based on IPSC amplitudes at 2 mM [Ca^2+^]_o_ in neurons of WT or *Scn1a*^+/–^ mice (WT: 1.36 ± 0.09 mM, *n* = 35, *Scn1a*^+/–^: 1.02 ± 0.04 mM, *n* = 36/*N* = 9 cultures, \*\*\**p* < 0.001).

### E/I balance of *Scn1a*^+/–^ neurons at high extracellular Ca^2+^ concentrations

Based on the high [Ca^2+^]_o_ sensitivity of the inhibitory synapses of *Scn1a*^+/–^ neurons, we hypothesized that *Scn1a* disruption might cause changes in the Ca^2+^ response of neurons. Because epileptic seizures are caused by excessive Ca^2+^ influx into nerve terminals^14^, we predicted that increasing [Ca^2+^]_o_ would induce a pseudo-epileptic condition and disrupt the E/I balance even in the normal state. We therefore investigated the magnification of excitatory and inhibitory synaptic transmission when [Ca^2+^]_o_ was changed from 2 mM to 8 mM. In excitatory neurons, there was no significant difference in the magnification of EPSCs between WT and *Scn1a*^+/–^ neurons (Figure 5c, d). This result is likely because EPSC amplitudes increase in proportion to an increase in [Ca^2+^]_o_ in an activity-dependent manner. Conversely, in inhibitory neurons, the IPSC amplitude magnification was significantly lower in *Scn1a*^+/–^ neurons than in WT neurons (Figure 5e, f). This finding indicates that IPSC amplitudes were unable to increase in proportion to the increase in [Ca^2+^]_o_. This phenomenon may be the result of significantly fewer synapses and increased [Ca^2+^]_o_ sensitivity at inhibitory synapses in *Scn1a*^+/–^ neurons. Combined with the finding of lower IPSC amplitude at 2 mM [Ca^2+^]_o_, these results indicate that, for *Scn1a*^+/–^ neurons, a higher [Ca^2+^]_o_ worsens the E/I balance.

## DISCUSSION

### E/I balance in the brains of *Scn1a*^+/–^ mice

Neurons in the brain form networks that function by maintaining a reciprocal E/I balance. When epilepsy develops, there is generally a temporary or persistent disruption of the E/I balance^13,14^. We therefore predicted that the E/I balance would be altered by either an increase in excitatory synaptic transmission or a decrease in inhibitory synaptic transmission. In the present study, we did not observe any enhancement of excitatory synaptic transmission at the single-neuron level in *Scn1a*^*+/–*^ neurons. However, inhibitory synaptic transmission was significantly lower in *Scn1a*^*+/–*^ neurons than in WT neurons. The lack of any differences in excitatory synaptic transmission may be because of the distribution of Na_v1.1_. These subunits have been reported to be expressed predominantly on the cell body of neurons^16,17^. Na_v1.1_ seems to be gradually expressed from postnatal day 10 (P10) and is substantially expressed after P18 in the total brain membrane fraction^8^. However, Na_v1.1_ is expressed at extremely low levels in excitatory neurons in the hippocampus^8^. Based on this finding, thalamocortical slices have previously been used from P14–20 of Na_v1.1_ mutant mice^18^. Moreover, the vulnerability of Na^+^ currents has been confirmed in inhibitory interneurons at P13–14 in a DS mouse model^7^. In culture conditions, the expression of Na_v1.1_ has been confirmed at 14 days *in vitro*^19^ and has even been noted at 7–9 days *in vitro*^20^. Thus, considering that the timepoint of Na_v1.1_ expression is not significantly different between *in vivo* and *in vitro* conditions, we speculated that Na_v1.1_ was likely to be expressed at 13–16 days *in vitro*, which was the timepoint that was used in our experiments. Although we did not confirm the expression of Na_v1.1_ in the mouse model used in the present study, it is possible that our electrophysiological results would support the previous finding that Na_v1.1_ is not expressed in hippocampal excitatory neurons. It has also been reported that *Scn1a*-deficient mice show abnormalities in inhibitory neuron function^7^, which is consistent with our results. However, in *in vivo* conditions, the disruption of *Scn1a* appears to cause abnormalities in both excitatory and inhibitory neurons^21^.

In contrast to the inhibition of inhibitory neurons, an increase in inhibitory autaptic current amplitudes in parvalbumin-positive interneurons has been reported in cortical slices from *Scn1a*^+/–^ animals^22^. In the present study, however, we used cultured autaptic neurons dissociated from the striatum of WT and *Scn1a*^+/–^ animals. Therefore, the effects of DS-related mutations on inhibitory synaptic transmission may differ between brain regions. In particular, as mentioned by De Stasi et al., the presence of multiple cells may cause some kind of homeostasis^22^. The neuronal network *in vivo* is complex, with multiple connections with various neurons, including not only excitatory and inhibitory neurons, but also catecholaminergic and cholinergic neurons. Therefore, autaptic currents onto parvalbumin-positive interneurons likely receive other synaptic inputs from surrounding neurons. However, our autaptic culture preparation was configured as a single neuron only, which allowed no synaptic input from other neurons. Considering that it has often been noted that the synaptic properties of autaptic single neurons are different from those of mass-cultured neurons^23,24,25^, future studies need to examine neural networks with mixed excitatory and inhibitory synaptic inputs.

### Synaptic function in *Scn1a*^+/–^ neurons

Although the RRP sizes at all synapses were similar between WT and *Scn1a*^+/–^ neurons, the number of GABAergic synapses was significantly lower in *Scn1a*^+/–^ neurons. These results suggest that the number of releasable synaptic vesicles per synapse is higher in *Scn1a*^+/–^ neurons than in WT. We had hypothesized that the lower P_vr_ in *Scn1a*^+/–^ neurons was caused by reduced sensitivity to [Ca^2+^]_o_; however, the results were contrary to our expectations (Figure 5b). Therefore, other mechanisms may affect the tethering system of Ca^2+^ influx into nerve terminals. For example, synaptotagmin is a Ca^2+^ sensor for synaptic release ^26^. It is thus possible that the function of synaptotagmin was affected in the nerve terminals of *Scn1a*^+/–^ neurons.

In general, epileptic seizures are caused by hyperexcitation of nerves, caused by the influx of large amounts of Ca^2+^ from extracellular sources. It has recently been demonstrated that a large amount of Ca^2+^ influx also occurs during epileptic seizures in animal models of DS^14^. In the present study, we simulated epileptic seizures by increasing the [Ca^2+^]_o_, and found that the E/I balance in *Scn1a*^+/–^ neurons was more disrupted during pseudo-epileptic seizures than during normal conditions. Although the mouse model used in this study only had mutated *Scn1a*, which encodes Na_v1.1_, there was an alteration in the Ca^2+^ response of *Scn1a*^+/–^ neurons. One reason for this may involve the expression of Ca^2+^ channels; it is known that *Scn1a* disruption alters the expression of Na^+^ channels as well as other ion channels^27^. Therefore, we speculate that disruption of *Scn1a* may cause changes in Ca^2+^ channels, thus leading to the alteration of Ca^2+^ responses in *Scn1a*^+/–^ neurons.

### Susceptibility of *Scn1a*^+/–^ mice to seizures

When epilepsy is induced, it may be triggered by a small external stimulus. For example, the firing of a spontaneous electrical potential, which generally has no physiological consequences, may stimulate epileptogenic cells to induce bursts of action potential firings, which then propagate through the neuronal network. Thus, even if a change in E/I balance does not have severe consequences in the normal condition, visible disruption may occur in the brains of *Scn1a*^+/–^ mice if they are experimentally subjected to a small number of external stimuli (e.g., frequent stimulation or drug stimulation).

A reduction in Na^+^ currents is the hallmark of DS, at least in juvenile mice. Notably, studies have demonstrated that a long-lasting depolarization elicits a vulnerability to Na^+^ currents^7,8,18,28^. These previous studies commonly reported that this vulnerability is observed within successive firings, indicating that long-lasting depolarization is required to observe a resulting decrease in Na^+^ currents. In the present study, we applied a short depolarizing pulse (2 ms) to elicit a synaptic response. As a result, the action potentials triggering the evoked synaptic transmission were identical between WT and *Scn1a*^+/–^ neurons of the hippocampus and striatum, respectively (Figure S2). We therefore hypothesized that a duration of 2 ms was too short to allow consecutive firings. However, given that epilepsy tends to occur in juveniles, additional external stimulation of juvenile synapses *in vitro* may be an effective way to detect small disruptions. At the cellular level, intracellular Cl^−^ concentrations are high in the juvenile stage and low in the mature stage^29,30,31^. In juveniles, we speculate that GABAergic input may cause Cl^−^ to leak out of cells via GABA_A_ receptors, resulting in cell depolarization. This developmental change may be one of the mechanisms that contribute to seizure susceptibility. However, a full validation of the molecular mechanisms requires further experiments.

In conclusion, the results of the present study suggest that mutations in the *Scn1a* gene can impair inhibitory synaptic transmission when extracellular Ca^2+^ concentrations are high.

## METHODS

### Vector construction

The structure of the targeting vector is shown in Figure 1. Briefly, a 5.4 kb murine *Scn1a* genomic fragment spanning from the end of exon 7 to the beginning of exon 5 (left arm) and a 5.0 kb murine *Scn1a* genomic fragment spanning from the end of exon 13 to the beginning of exon 16 (right arm) were amplified by polymerase chain reaction (PCR) from the bacterial artificial chromosome (BAC) clone TRPCI23-201O18 (Advanced GenoTechs). The accession number of the corresponding *Scn1a* gene reference sequence is NC_000068.7. The resulting construct was modified to contain a lox71 sequence downstream of *Scn1a* exon 7, and a neomycin-resistance (neo) cassette was added further downstream, which comprised PPGK (the promoter sequence of the mouse phosphoglycerate kinase 1 [*Pgk*] gene), a lox2272 sequence, and a *Pgk* polyadenylation sequence. The neo cassette was additionally flanked by two yeast-derived flippase (FLP) recognition target sequences (FRTs) to allow the removal of the drug-resistance cassette in a flippase-enabled background. The *Scn1a* fragment was 5′-fused with a diphtheria toxin-A cassette, PCR subcloned from plasmid p03. The full construct was inserted into the multiple cloning site of pBluescript II SK (+) (Stratagene).

### Culture and gene manipulation of mouse embryonic stem (ES) cells

Feeder-free mouse ES cells, which were TT2-KTPU8 strain (C57BL/6J × CBA) F1-derived wild-type ES cells, were seeded at 6.0 × 10^6^ cells per dish in collagen-coated 10-cm diameter dishes containing Glasgow minimum essential medium (Thermo Fisher Scientific) supplemented with 1% fetal calf serum, 14% knock-out serum replacement (Thermo Fisher Scientific), and 1 kU/mL leukemia inhibitory factor (Thermo Fisher Scientific). After a 2-day incubation at 37°C in 6.5% CO_2_ to reach semi-confluence (1.6 × 10^7^), the cells were washed twice with phosphate-buffered saline, resuspended in 1.6 mL phosphate-buffered saline, and divided into two 0.8-mL aliquots. Transfections were performed using a Gene Pulser (Bio-Rad) and a 4-mm gap cuvette set at 0.8 kV/3 µF in the presence of a 20 µg PvuI-linearized targeting vector. Transfected cells were expanded in G418-supplemented medium (200 µg/mL) for 9 days. Cells with randomly inserted target vector DNA did not survive during this time because non-homologous recombination failed to remove the diphtheria toxin-A cassette. Of the approximately 500 colonies that developed, 192 were isolated by pipetting under a microscope and were then cultured for 3 days. Next, two-thirds of these cells were suspended in dimethyl sulfoxide-supplemented culture medium and cryo-banked. The remaining cells were used for genomic DNA extraction, followed by PCR and Southern analysis. The PCR primers RA/S and RA/A, which were used to amplify a 0.6-kb DNA fragment of hybridization probe, were designed according to the nucleotide sequences of intron 13 and exon 13 of *Scn1a*, respectively (Figure 1b). The resulting DNA fragments were used as the probe for Southern blot analysis following EcoRI-/PvuI-digestion of genomic DNA, which hybridized with 7.8 kb and 6.5 kb fragments for the non-recombinant and recombinant alleles, respectively (Figure 1c).

### Animals

Experimental animals were handled under the ethical regulations for animal experiments of the Fukuoka University experimental animal care and use committee and animal experiment implementation. All animal protocols were approved by the ethics committee of Fukuoka University (permit numbers: 1712128 and 1812092). All experimental protocols were performed according to the relevant guidelines and regulations by Fukuoka University. All *in vivo* work was carried out in compliance to ARRIVE guidelines.

Pregnant ICR mice for the astrocyte (a type of glial cell) cultures were purchased from CLEA Japan, Inc. (Catalogue ID: Jcl:ICR). Mice were individually housed in plastic cages and kept in an animal house under the following conditions: room temperature 23°C ± 2°C, humidity 60% ± 2%, and a 12-hour light/dark cycle (7:00 AM lights on), with free access to water and food (CE-2, Nihon Crea).

### Development of *Scn1a*^+/–^ mice

Gene targeted clones of *Scn1a*-mutant ES cells were used to generate *Scn1a*-812neo mice. We first removed two-cell stage zona pellucida cells from the embryos of the Institute for Cancer Research (ICR) mouse line and immersed them in mutant ES cell suspension. After an overnight incubation to enhance the aggregation of embryos and cells, chimeric embryos were transplanted into the uteri of pseudopregnant females. From the resulting litter, checker-coated chimeras were mated with C57BL/6J mice to produce F1 heterozygous offspring with a single mutant *Scn1a* allele. The resulting mouse line containing the *Scn1a* mutation was backcrossed with C57BL/6J mice over 10 generations to establish a congenic strain. Mice with an *Scn1a* gene in which exon 8 to exon 12 was heterogeneously replaced with a neomycin resistance gene were termed *Scn1a*^+/–^ mice.

### Genotyping

Genomic DNA was extracted from mouse tails by NaOH extraction. The PCR primers Scn1ai7prS2 and FRT-RV, which were used to detect the recombinant allele, were designed according to the nucleotide sequences of intron 7 of the *Scn1a* gene and FRT elements, respectively. Using PCR, these primers only amplified a 0.6-kb DNA fragment when genomic DNA containing the homologous recombination was used as the template. Newborn mice were used for the functional analysis of primary neuronal cultures after PCR verification.

### Microisland cultures of astrocytes

We used a “coating stamp” to obtain uniformly sized cell adhesion areas, divided into dots (squares with a side length of about 300 μm)^32,33^. The coating stamp made it possible to minimize variations in the size of each island of astrocyte culture^32,33^.

ICR mice immediately after birth (P0–1) were used for astrocyte cultures. Details of the astrocyte culture method are described in our previous paper^32,33^.

### Autapse culture specimens of single neurons

We used autapse culture preparations, which are co-cultured preparations of dot-cultured astrocytes and a single neuron, for the synaptic function analysis^32,33,34,35,36^. The axons of autapse neurons form many synapses (autapses) to their cell bodies and dendrites, without any synaptic contacts from other neurons. This feature allowed us to focus on the synapses of a single neuron only, and to analyze the properties of synaptic function in more detail.

*Scn1a* knock-in neurons (*Scn1a*^+/–^ neurons) were used in all experiments. For the culture of single neurons, *Scn1a*^+/–^ mice immediately after birth (P0–1) were decapitated to minimize pain/discomfort and used; tails were harvested to confirm disruption of the *Scn1a* gene, and PCR was used for genotyping. Neurons were enzymatically isolated from the hippocampus of P0–1 *Scn1a*^+/–^ mice using papain (Worthington). Before plating neurons, the conditioned medium of the microisland cultures was replaced with Neurobasal A medium (Thermo Fisher Scientific) containing 2% B27 supplement (Thermo Fisher Scientific) and 1% GlutaMAXLI supplement (Thermo Fisher Scientific). Cells were plated at a density of 1500 cells/cm^2^ onto the astrocyte microislands. Hippocampal neurons were used to analyze excitatory synaptic transmission and striatal neurons were used to analyze inhibitory synaptic transmission. To maintain the soluble factors from astrocytes, the medium was not changed during the culture period. The incubation period was 13–16 days *in vitro*.

### Electrophysiology

Synaptic analysis was performed using the patch-clamp method, where the membrane potential was clamped at –70 mV. An evoked excitatory postsynaptic current (EPSC) or inhibitory postsynaptic current (IPSC) was recorded in response to an action potential elicited by a brief (2 ms) somatic depolarization pulse (to 0 mV) from the patch pipette under voltage-clamp conditions. Action potentials triggering EPSCs or IPSCs were not significantly different between wild-type (WT) and *Scn1a*^+/–^ neurons of the hippocampus and striatum, respectively (Figure S2). Spontaneous miniature EPSCs (mEPSCs) and IPSCs (mIPSCs) were recorded in the presence of a Na^+^ channel inhibitor, 0.5 μM tetrodotoxin (FUJIFILM Wako Pure Chemical Corporation), in the extracellular fluid. The readily releasable pool (RRP) size was calculated as the total area of inward current elicited by transient hyperosmotic extracellular fluid containing 0.5 M sucrose. The vesicular release probability (P_vr_) was calculated by dividing the evoked EPSC or IPSC area by the RRP area. Synaptic responses were recorded at a sampling rate of 20 kHz and were filtered at 10 kHz. Data were excluded from the analysis if a leak current of > 300 pA was observed. The data were analyzed offline using AxoGraph X 1.2 software (AxoGraph Scientific).

Extracellular fluid (pH 7.4) was as follows (mM): NaCl 140, KCl 2.4, HEPES 10, glucose 10, CaCl_2_ 2, and MgCl_2_ 1. Intracellular fluid (pH 7.4) for recording EPSCs was as follows (mM): K-gluconate 146.3, MgCl_2_ 0.6, ATP-Na_2_ 4, GTP-Na_2_ 0.3, creatine phosphokinase 50 U/mL, phosphocreatine 12, EGTA 1, and HEPES 17.8. The intracellular fluid for recording IPSCs was prepared by replacing K-gluconate with KCl in the intracellular fluid for EPSCs. All experiments were performed at room temperature (20°C –22°C).

### Immunocytochemistry

Striatal neurons cultured for 14 days were fixed by being placed in 4% paraformaldehyde in phosphate-buffered saline for 20 minutes. They were then incubated with primary antibodies containing blocking solution (0.1% Triton X-100, 2% normal goat serum) at 4°C overnight. The following primary antibodies were used: anti-microtubule-associated protein 2 (MAP2 (Cat. # 188 004); guinea-pig polyclonal, antiserum, 1:1000 dilution, Synaptic Systems) and anti-vesicular GABA transporter (VGAT (Cat. # 131 002); rabbit polyclonal, affinity-purified, 1:2000 dilution, Synaptic Systems). Next, the cells were incubated in secondary antibodies corresponding to each primary antibody (Alexa Fluor 488 for MAP2 and Alexa Fluor 594 for VGAT, 1:400, Thermo Fisher Scientific) for 40 minutes at room temperature. The nuclei of astrocytes and autaptic neurons were visualized by counterstaining with 4’,6-diamidino-2-phenylindole (DAPI)-containing mounting medium (ProLong® Gold Antifade Reagent with DAPI, Thermo Fisher Scientific). Only neurons with one nucleus were analyzed when counting synapse numbers.

### Image acquisition and quantification

Confocal images of autaptic neurons were obtained using a confocal microscope (LMS710, Carl Zeiss) with a C-Apochromat objective lens (40×, NA 1.2, Carl Zeiss), with sequential acquisition at high-resolution settings (1024 × 1024 pixels). The parameters of each image were optimized for the z-stack setting (a depth of 9 μm with 0.5 μm steps) and pinhole setting (1 Airy unit, 0.9 μm). The depth sufficiently covered the thickness of neurites in our autaptic culture. The confocal microscope settings were identical for all scans in each experiment. A single autaptic neuron was selected for each scan in a blinded manner, based on MAP2 fluorescence images. The sizes and numbers of synaptic puncta were detected using ImageJ software. The procedures used to analyze the synaptic puncta have been published previously^36,37^.

### Statistical analysis

All data are expressed as the mean ± standard error. A lowercase *n* indicates the number of neurons recorded, and an uppercase *N* indicates the number of cultures (lot number). Two groups were compared using the Student’s unpaired *t*-test. Statistical significance was considered when *p* < 0.05.

### Data accessibility statement

The data that support the findings of this study are available from the corresponding author on reasonable request.

## ACKNOWLEDGMENTS

This work was supported by a KAKENHI Grant-in-Aid for Scientific Research (C) to S.K. (No. 17K08328) from the Japan Society for the Promotion of Science, and the Science Research Promotion Fund and The Fukuoka University Fund to S.H. (Nos. G19001 and G20001), a grant for Practical Research Project for Rare/Intractable Diseases from Japan Agency for Medical Research and development (AMED) to S.H. (Nos. 15ek0109038h0002 and 16ek0109038h0003), a KAKENHI Grant-in-Aid for Scientific Research (A) to S.H. (No. 15H02548), a KAKENHI Grant-in-Aid for Scientific Research (B) to S.H. (Nos. 20H03651, 20H03443 and 20H04506), the Acceleration Program for Intractable Diseases Research utilizing Disease-specific iPS cells from AMED to S.H. (Nos. 17bm0804014h0001, 18bm0804014h0002, and 19bm0804014h0003), a Grant-in-Aid for the Research on Measures for Intractable Diseases to S.H. (H31-Nanji-Ippan-010), Program for the Strategic Research Foundation at Private Universities 2013-2017 from the Ministry of Education, Culture, Sports, Science, and Technology (MEXT) to S.H. (No. 924), Center for Clinical and Translational Research of Kyushu University Hospital to S.H. (No. 201m0203009 j0004). We thank the members of our laboratory for their help. We also thank Bronwen Gardner, PhD, from Edanz Group (https://en-author-services.edanz.com/ac) for editing a draft of this manuscript.

## AUTHOR CONTRIBUTIONS STATEMENT

K.U., H.K., Y.T., Y.A., and A. A. performed experiments and analyzed data; Y.T and M.D. created the *Scn1a*^+/–^ mouse model; S.K. conceived the study; K.K., T.W., K.I., and S.H. interpreted the data; K.U. and S.K. wrote the manuscript with input from all authors. All authors reviewed the manuscript.

## ADDITIONAL INFORMATION

### Competing interests

None of the authors has any conflict of interest to disclose.

### Ethical publication statement

We confirm that we have read the Journal’s position on issues involved in ethical publication and affirm that this report is consistent with those guidelines.

## FIGURE LEGENDS

**Figure S1**. Raw data underlying Figure 1c.

(**a**) Full-length image of the Southern hybridization for seven candidate clones. Handwritten characters indicate the identification numbers of the samples.

**Figure S2**. No changes in Na^+^ current amplitudes in wild-type (WT) and *Scn1a*^+/–^ neurons of the hippocampus and striatum.

(**a**) Representative traces of Na^+^ currents evoked by depolarizing pulses (–70 mV to 0 mV) via patch pipette for 2 ms under voltage-clamp conditions. There were no differences in Na^+^ currents between the WT and *Scn1a*^+/–^ neurons of the hippocampus and striatum. (**b**) Average amplitudes of the Na^+^ current in WT and *Scn1a*^+/–^ neurons of the hippocampus and striatum.

## Notes

### Competing Interest Statement

The authors have declared no competing interest.

### Summary of Updates

The title has been changed to make more straightforward and precise scientific meaning.

