## Supplemental Figures for "Inhibitory synaptic transmission is impaired at higher extracellular Ca^2+^ concentrations in *Scn1a*^+/−^ mouse model of Dravet syndrome"

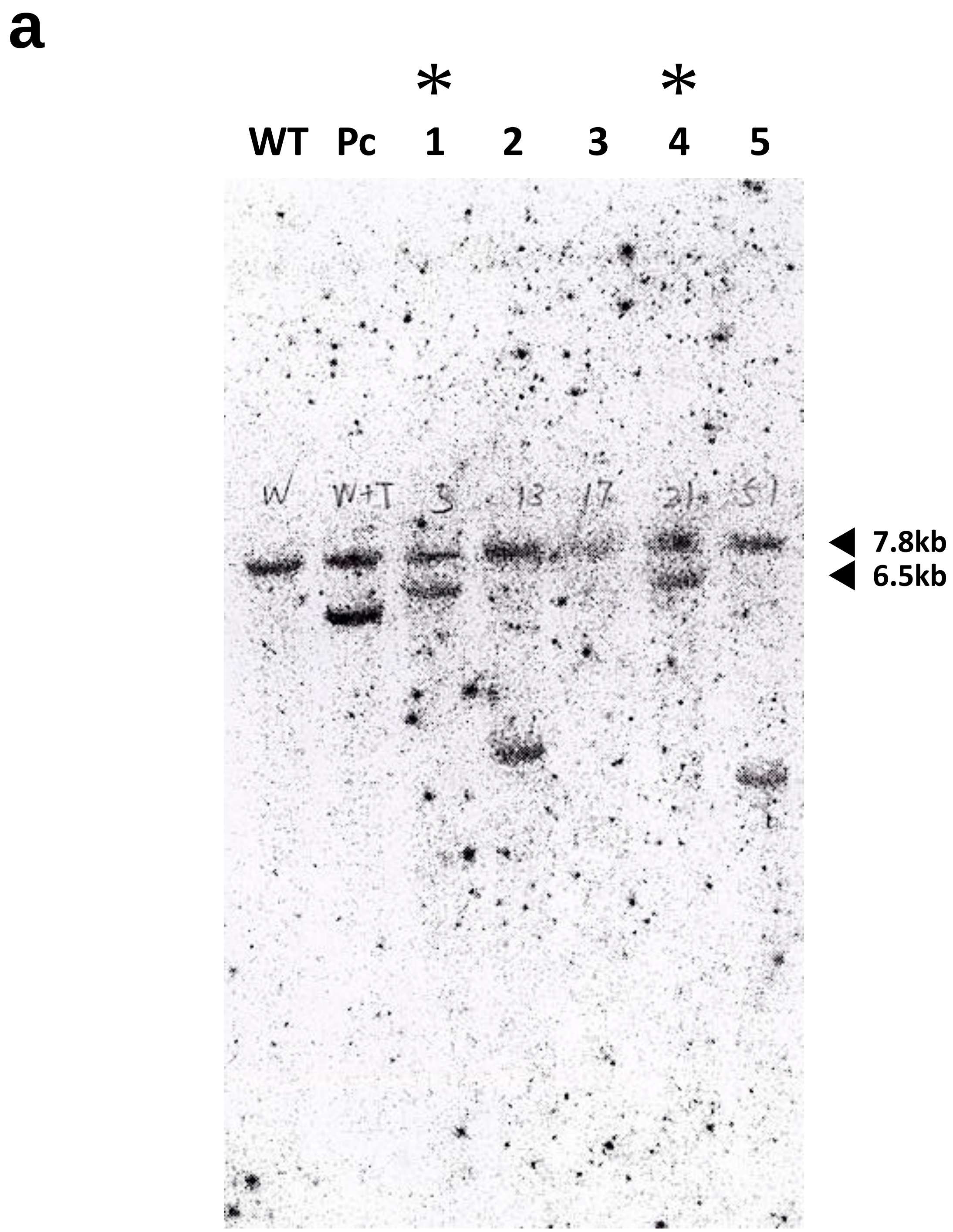


**Figure S1.** Raw data underlying Figure 1c.
(**a**) Full-length image of the Southern hybridization for seven candidate clones. Handwritten characters indicate the identification numbers of the samples.

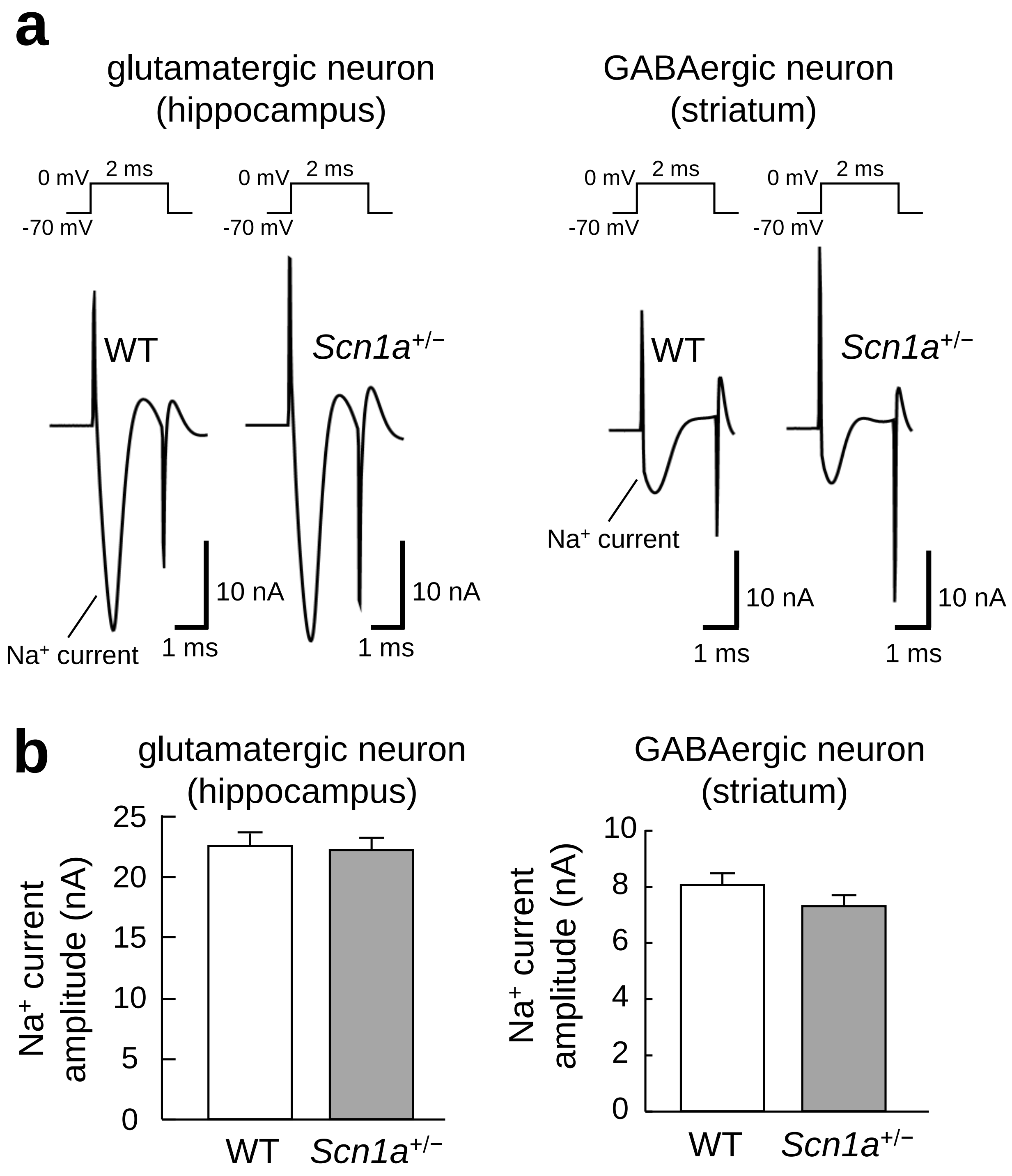


**Figure S2.** No change in the $Na^+$ current amplitude in WT and *Scn1a*$^{+/-}$ neurons of the hippocampus and striatum.

(**a**) Representative traces of $Na^+$ currents evoked by depolarizing pulses (-70 mV to 0 mV) via patch pipette for 2 ms under the voltage-clamp condition. Note that there is no difference in $Na^+$ currents between WT and *Scn1a*$^{+/-}$ neurons of the hippocampus and striatum. (**b**) Average amplitudes of the $Na^+$ current in WT and *Scn1a*$^{+/-}$ neurons of the hippocampus and striatum.
